# Fear engrams and NPYergic circuit in the dorsal dentate gyrus determine remote fear memory generalization

**DOI:** 10.1101/2022.04.26.489543

**Authors:** Syed Ahsan Raza, Katharina Klinger, Miguel del Ángel, Yunus Emre Demiray, Gürsel Çalışkan, Michael R. Kreutz, Oliver Stork

**Author notes:** These authors contributed equally to this work (Katharina Klinger, Miguel del Ángel). Corresponding authors, and, Address: Department of Genetics and Molecular Neurobiology, Institute of Biology, Otto-von-Guericke University Magdeburg, Leipziger Str. 44, Haus 91, 39120 Magdeburg, Germany.

## Abstract

Generalization is a critical feature of aversive memories and significantly contributes to post-traumatic stress disorder (PTSD) pathogenesis. While fear memories over time tend to generalize across differences in the contextual background and even to novel contextual settings, this effect can be counteracted by exposure to controlled reminder sessions even at remote time points. Using Pavlovian fear conditioning in mice, we show that generalization to a novel context of remote memory is associated with a loss of cellular engram activation in the dorsal dentate gyrus (dDG) and can be effectively counteracted by a preceding contextual reminder session. In addition to engram cells activation in response to a novel context, the reminder session also leads to a recovery of neuropeptide Y (NPY) function in the dDG and dDG-CA3 neurotransmission. In line with a proposed role of NPY as a resilience factor, we found that chronic viral knockdown of NPY in the dDG and blockage of its activity-dependent expression in NPYergic dDG interneurons with dominant-negative CREB^S133A^ both increase remote memory generalization. With chemogenetic silencing of these interneurons, we could localize their critical involvement to a time window during and immediately following the fear memory acquisition. Together, these findings suggest that NPYergic interneurons of the dDG, shaping the memory engram during fear learning and early consolidation, determine fear generalization.

## Introduction

The generalization of learned fear is an adaptive coping mechanism; it allows animals to quickly adjust their behavior to novel situations based on their experience made in an ever-changing environment. However, over-generalization can occur after traumatic events and is typically observed in posttraumatic stress disorder (PTSD) and generalized anxiety disorders (1–3). It can lead to inappropriate anxiety (4) and disturb the ability to discriminate threats from safety (5). Strikingly, remote memories are more generalized than recent ones, which suggests a time-dependent reorganization in hippocampal-cortical networks (6–10). Therefore, a better understanding of such remote memory generalization mechanisms may thus guide the search for an effective new intervention strategies.

Pavlovian fear conditioning in rodents is widely used as a tool to model PTSD and understand the involved limbic lobe circuits of fear generalization (11–17). The dorsal dentate gyrus (dDG) has emerged as a critical structure during learning that hosts part of the cellular engram for context memory (18–19). The dDG controls the specificity and generalization of contextual fear memories through its pattern separation and pattern completion functions (20–21). Indeed, accumulating evidence suggests that the dDG is critically involved in the contextual generalization of remote memories (22–24).

Activity in the DG is tightly controlled by local GABAergic circuits (25), amongst which the hilar inhibitory perforant-path (HIPP) cells may be of particular interest. We and others have previously shown that the HIPP cells control the encoding of fear memory salience in the DG engram (26–27). The HIPP cells co-release Neuropeptide Y (NPY) and somatostatin (SST) to inhibit granule cells as well as other hilar interneurons (28). We have previously demonstrated that NPY mediates context fear salience encoding via postsynaptic NPY-Y1 receptors in the dDG (27). Moreover, NPY is implicated as a resilience factor in PTSD (29–32), and in fact, we have demonstrated that genetic suppression of NPY expression in the dDG increases the proportion of affected animals in a mouse model of PTSD (33)

In the current study, we investigate the beneficial effect of context memory recall in reducing the generalization of fear memories to novel situations. We demonstrate with engram labeling and slice physiology that fear recall results in a reactivation of cellular responsiveness to novel context settings and perforant path-induced activation in the dDG-CA3 circuit. Moreover, with targeted viral intervention, we demonstrate the specific role of NPYergic HIPP cells in these mechanisms.

## Materials and Methods

### Animals

Wild type C57BL/6BomTac mice (M&B Taconic, Germany) and StatinCre (Sst^tm2.1(cre)Zjh^/J) were raised and bred with *ad libitum* food and water in the animal facility at the Institute of Biology, Otto-von-Guericke University Magdeburg. Adult male mice (10-12 weeks) were housed in groups of 2-6 on an inverse 12 h light/dark cycle (lights on at 7 pm with a 30 min dawn phase). Before starting the behavioral experiments, mice are single caged for 2-3 days (otherwise mentioned). Littermates were randomly assigned to experimental and control groups and tested in parallel. Animal housing and animal experiments were in accordance to the European and German regulations for animal experiments and approved by the Landesverwaltungsamt Saxony-Anhalt (Permission Nr. 42502-2-1563-UniMD).

### Fear conditioning

Single caged mice were handled for two days before the start of the experiment. All experiments were performed during the dark phase. We used cued conditioning protocol as described earlier (27). Mice were conditioned in a soundproof chamber containing a 16 cm × 32 cm × 20 cm acrylic glass arena with an attached grid floor to deliver foot shocks, a loudspeaker, and an exhaust fan (background noise 70 dB SLP, light intensity < 10 lux; TSE Systems). Mice were first acclimatized to the conditioning chamber for 6 min each day for two days to provide a pre-training representation of contextual features. After 120 s of contextual exposure on the third day, paired fear conditioning was performed with the presentation of three conditional stimuli (CS: 10 kHz tone for 10 s, 80 dB), each co-terminating with an unconditional stimulus (US foot shock: 0.5 mA for 1 s), with inter-stimulus intervals (ISI) of 20 s. In unpaired conditioning, 3 US and 3 CS are presented without temporal coincidence with variable ISIs ranging from 20-40 s. Mice are placed in their home cages 120 s after the last stimulus. During acclimatization and training, chambers were cleaned and scented with 70% ethanol, providing additional sensory cues. After training, fear recall is tested either at 24 h (recent memory) or five weeks (remote memory) time points. As a novel-context, we used clean standard cages without bedding material, striped with black stickers from outside, cleaned and scented with 2% acetic acid. Except for the background noise, this arrangement minimizes the similarities to the conditioning chamber. For recall, mice were placed in the novel-context for 240 s without any shock and CS. The recall in the shock-context was tested in the original conditioning chamber scented with 70% ethanol 2 h after the novel-context exposure. After the initial 240 s, 4 CS tones were played to test auditory memory only in the shock-context. The animals’ freezing behavior was evaluated online via a photobeam detection system that detected immobility periods >1 s. The freezing score was calculated as a percentage (+ SEM) of total time spent freezing (two min bouts) during context exposure and separately during CS presentation.

### Viral vectors and Drugs

Adeno-associated viruses (AAV), AAV-hSyn-DIO-hM4Di-mCherry, AAV-hSyn-DIO-hM3Dq-mCherry, and AAV-hSyn-DIO-mCherry deposited by Dr. Roth were purchased from Addgene (USA). Lentiviruses, LV-shNPY-EGFP, and LV-2xfl-CREB^S133A^-HA were designed and prepared in our lab (27, 33). The pAAV-RAM-d2TTA::TRE-MCS-WPRE-pA plasmid deposited by Dr. Lin was purchased from Addgene (USA) and AAV-RAM particles were prepared at the UKE vector facility with a 4.25 × 10^13^ vg/ml titer. DREADD receptors were activated 1 h before the respective task via i.p. injection of 10 mg/kg clozapine-N-oxide (CNO) (Hellobio, UK) in physiological saline.

### Stereotaxic surgery

Mice were anesthetized with 5% isofluorane in O2/N2O mixture (combi-vet® Rothacher Medical GmbH, Switzerland), and anesthesia was maintained at 1.5-2% mixture during the surgery. After placing on a stereotaxic frame (World Precision Instruments, Germany), craniotomy was performed targeting the dDG (Paxinos & Watson brain atlas for mouse) anterioposterior (AP): −1.94 mm, mediolateral (ML): ±1.3 mm from Bregma and dorsoventral (DV): −1.7 mm from the brain surface. Viral vectors were injected with a digital microsyringe pump (World Precision Instruments, Germany) at a flow rate of 0.1 μl per min through a 33G injection needle fitted in a 10 μl NanoFil microsyringe. Each mouse was injected subcutaneously with 5 mg kg^−1^ carprofen to linger pain during surgery. Mice were allowed 7-10 days for virus expression and recovery from surgery until the start of fear conditioning. Each injection was made with 1 μl of virus with following titer, AAV-hM4Di 1.8 × 10^13^, AAV-hM3Dq 3.2 × 10^13^, AAV-mCherry 2.2 × 10^13^, LV-shNPY 1.1 × 10^8^ and LV-CREB^S133A^ at 7.8 × 10^9^.

For the engram labeling experiment, 500 nl per area AAV-RAM viral vectors at 1:14 dilution in sterile PBS are injected two days after mice were fed with doxycycline chow (40 mg kg^−1^). The labeling window was generated by replacing the doxycycline chow immediately after the last acclimatization phase with standard chow. In this experiment, training was performed 48 h after the last acclimatization. After training, mice are placed back on doxycycline chow till sacrificed.

### Immunohistochemistry

Immunohistochemistry was performed as described earlier (27) on 30 μm thick coronal sections obtained from brains perfused with 4% PFA. For cFos labeling, mice were perfused 90 min after the contextual exposure. For NPY labeling, mice were perfused 24 h or 5 weeks after training without contextual exposure. Anti-cFos (Cell Signaling #2250) at 1:1000, anti-NPY (GTX #10980) at 1:10,000, anti-HA (BioLegend) at 1:300 were used as primary antibodies, and biotinylated goat anti-rabbit at 1:200 (Vector Laboratories, USA) as a secondary antibody, which is visualized with Cy5 (1:1000, Jackson ImmunoResearch Labs, UK). Leica DMi8 Fluorescent microscope (Leica, Germany) was used to perform imaging. Area sizes in the dDG granule cell layer for cFos, and dDG hilus for NPY were marked with ImageJ (NIH), and cFos^+^ cells and NPY^+^ cells were manually counted, respectively. Cell densities were expressed as the number of positive cells per mm^2^. We used mean cell densities from 6 sections per animal for statistical comparison between groups.

For quantification of ensemble reactivation, mean cell densities for RAM^+^, cFos^+^, and double^+^ (RAM^+^-cFos^+^) cells were used from 6 sections per animal. The percent reactivation rate is calculated as (# of double^+^ cells / RAM^+^ x 100).

### Peptide isolation and ELISA

Mice were killed by cervical dislocation either 24 h or 5 weeks after training without contextual exposure, and brains were carefully removed from the skull and placed in ice-cold PBS. The left and right brain hemispheres were separated along the longitudinal fissure with a scalpel, and then the thalamus and hypothalamus were removed with forceps to expose the hippocampus. The dorsal part of the DG was carefully removed from CA1 and CA3 boundaries (34) and stored at −80°C. Stored DG tissues were homogenized in cold LM Buffer (1% Laurylmaltoside, 1% NP-40, 1mM Na3 VO4, 2mM EDTA 50 mM Tris-HCl pH 8.0, 150 mM NaCl, 0.5% DOC, 1mM AEBSF, 1uM Pepstatin A, 1mM NaF, 1 Tablet of Pierce protease inhibitor.) 100 ul of the protein extract was used in each sample to determine NPY peptide levels via Enzyme-linked Immunosorbent Assay (ELISA) according to manufacturer’s instructions (Uscn Life Science).

### Field potential recordings

Mice were deeply anesthetized with isoflurane and decapitated. Brains were rapidly removed and placed into cold (4–8 °C) carbogenated (5% CO_2_/95% O_2_) artificial cerebrospinal fluid (aCSF) containing (in mM) 129 NaCl, 21 NaHCO_3_, 3 KCl, 1.6 CaCl_2_, 1.8 MgSO_4_, 1.25 NaH2PO_4_, and 10 glucose. Parasagittal slices containing the dorsal hippocampus were obtained by cutting the brain at an angle of about 12° on an angled platform. The four most dorsal slices were transferred to an interface chamber perfused with aCSF at 32 ± 0.5 °C (flow rate: 1.8 ml ± 0.2 ml per min, pH 7.4, osmolarity ~ 300 mOsmol kg^−1^). After cutting, the slices were left for recovery for at least 1 h before starting recordings. The perforant path (PP) stimulation was performed with a bipolar tungsten wire electrode, with exposed tips of ~ 20 μm and tip separations of ~ 75 μm (electrode resistance in aCSF: ~ 0.1 MΩ, World Precision Instruments, Berlin, Germany). A glass electrode filled with aCSF (~ 1 MΩ) was placed at 70-100 μm depth into the granule cell layer (GL) of dDG to measure the population spike (PS) and another electrode in the stratum lucidum (SL) of dCA3 to measure the mossy fiber (MF)-mediated disynaptic field excitatory postsynaptic potential (fEPSP) response (35). An input/output (I/O) curve was recorded after stable responses were established (0.033 Hz, pulse duration: 100 μs, for 20 min). Pulses with intensities ranging from 10 to 50 μA were applied to obtain the I/O curve. The stimulus intensity that resulted in ~ 70% of the maximum PS amplitude in the dDG was further used to assess the effects of NPY Y1 receptor blockade on PS in the dDG and fEPSP responses in the dCA3. After 20 min of baseline recording, the selective Y1 receptor antagonist BIBP3226 (1 μM) was added to the aCSF for 30 min (0.033 Hz, pulse duration: 100 μs). Signals were pre-amplified using a custom-made amplifier and low-pass filtered at 3 kHz. Signals were sampled at a frequency of 10 kHz and stored on a computer hard disc for offline analysis (Spike2 software version 8; Cambridge Electronic Design, Cambridge, UK). The amplitudes of PS (average of ascending and descending component of PS) and disynaptic fEPSP (longer-latency descending component) were calculated using self-written MATLAB-based analysis tools (MathWorks, Natick, MA, USA) and the Spike2 software.

### Statistical analysis

Data are shown as mean + SEM, and each figure legend contains *n* numbers used for statistical analysis. Data analysis was done with GraphPad Prism, and *p* values are reported in the corresponding figure legends. Briefly, Student’s two-tailed *t*-test was used to compare two groups and paired *t*-test or Wilcoxon signed-rank test was used to compare matched datasets. One-way analysis of variance (ANOVA) and two-way ANOVA were used for multiple comparisons (repeated measures for matched datasets), followed by Fisher’s protected least significant difference test or Tukey’s multiple comparison test for post-hoc comparisons.

## Results

### Recall of fear 5 weeks after conditioning results in remote memory generalization

We first established a fear-conditioning paradigm to study recent and remote memory in C57BL/6 mice (Fig. 1a). Mice conditioned with a paired presentation of tone and shock showed significantly higher freezing upon recall in the novel-context after five weeks (remote) of fear incubation compared to one day (recent) after conditioning (*t*_14_ = 4.60, *p* = 0.0005). These groups showed no difference when tested 120 min later in the shock-context and during conditional stimulus (CS) tones (Fig. 1b). The baseline and post-shock freezing were not different between groups (Supplementary Fig. 1a). We found similar remote fear generalization in mice with unpaired conditioning protocol (Supplementary Fig. 1b) and in mice that remained group-housed till remote recall (Supplementary Fig. 1c). This data indicates a time-dependent increase in fear memory generalization after five weeks of fear incubation.

**Fig. 1:**
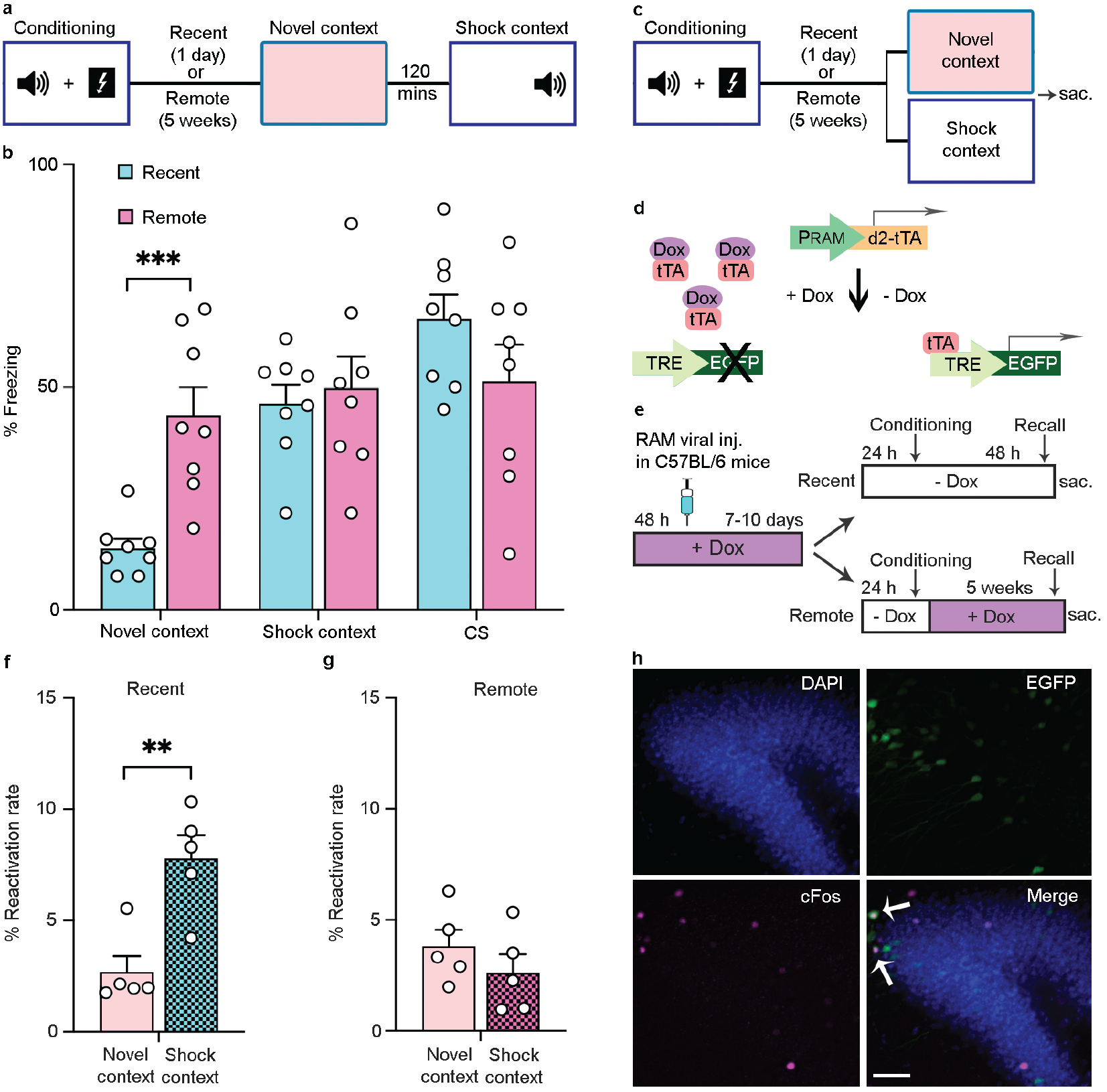
Remote fear memory generalization and recruitment of fear memory engrams in the dDG. **a** Schematic of the fear generalization paradigm. **b** Recall of fear in the novel context five weeks after the fear memory training results in significantly higher freezing behavior than the recall after one day (*n* = 8 each). Freezing behavior is comparable between the recent and remote groups in the shock-context and during CS presentation. **c** Schematic overview of the paradigm for engram labeling. **d** Schematic of the robust activity marking (RAM) system for engram labeling. tTA, tetracyclin transactivator, Dox, doxycycline. **e** Scheme of the engram labeling experiment. Mice received an AAV-RAM-EGFP (RAM) injection and acclimatization while on doxycycline chow (+DOX). 24 h after a change to standard chow (-DOX), mice were fear conditioned. For recent memory, mice remained on standard chow, and recall was tested 48 h later. For remote memory, mice were returned to +DOX chow, and recall was tested five weeks later. Tissue was collected 90 min after the respective test session. **f** In recent memory, shock-context exposure results in a significantly higher overlap of RAM^+^ and cFos^+^ (% reactivation rate) cells in the dDG than novel-context exposure (*n* = 5). **g** By contrast, the % reactivation rate remains low after both remote novel- and remote shock-context exposure (*n* = 5). **h** Representative microscopic images of the dDG show DAPI (blue), RAM-EGFP^+^ (green), cFos^+^ (magenta). Arrows depict double^+^ cells in a merged image. Scale bar, 100 μm. Data are mean + SEM. Statistics were performed using Student’s two-tailed *t*-test. \*\*\**p*<0.001, \*\**p*<0.005.

### Remote fear incubation decreases dDG engram size and results in fear generalization

We next used cFos-based analysis of neuronal activation patterns to study the ensembles of remote fear generalization in the dDG. We used cFos-dependent Robust Activity Marking (RAM) reporter system to monitor engram activation in the dDG. This doxycycline-dependent Tet-Off system labels the active neurons during conditioning (RAM^+^), and cFos^+^ labeling is used for fear recall in the same animal. The co-labeling of RAM^+^/cFos^+^ thus indicates reactivated engram populations (Fig. 1c-e) (36). In recent recall groups, we found a significant increase in engram labeling in the dDG after shock-context compared to the novel-context exposure (Fig. 1f and h, *t*_8_ = 4.06, *p* = 0.004). Intriguingly, this difference was lost in remote recall groups, where the engram labeling in the shock-context was reduced in size and similar to the novel-context group (Fig. 1g-h). In all groups, the RAM^+^ labeling and cFos^+^ ensemble size of the dDG after conditioning remained unchanged (Supplementary Fig. 2a-f). It has been shown that the cFos ensemble size in the anterior cingulate cortex is significantly larger after remote memory generalization (37–38), which we also confirmed in our experiment (Supplementary Fig. 2g-h). Our data suggest that the alteration in the dDG fear-engram activity over time is a cellular correlate for a time-dependent increase in fear generalization.

### Remote memory contextual reminder reinstates the dDG engram activation and attenuates fear generalization

Contextual re-exposure allows re-consolidation of the original memory and, in return, decreases its generalization (39–43). Importantly, mild exposure to the fear cues is used to weaken fear salience value during exposure therapy in human PTSD patients (44–46). Therefore, we hypothesized that a brief contextual reminder before remote recall would possibly decrease generalization in the novel-context by reactivating the engram ensembles in the dDG. Indeed, a contextual reminder session four weeks after conditioning significantly decreased freezing in the novel-context tested one week later while freezing during the recall in the shock-context and during tones remained unchanged (Fig. 2a-b, *F*_2.22_ = 11.13, *p* < 0.001; post hoc: recent vs. remote: *p* < 0.001; remote-reminder vs. remote: *p* = 0.006). The baseline and post-shock freezing were not different between groups (Supplementary Fig. 3a). Moreover, the reminder session reinstated activation of the dDG engram activity to levels found after recent memory, as indicated by RAM engram/cFos labeling (Fig. 2c-d, *F*_2.12_ = 16.52, *p* < 0.001; post hoc: recent vs. remote: *p* = 0.002; remote-reminder vs. remote: *p* < 0.001) without affecting either the total RAM or the total cFos^+^ labeling (Supplementary Fig. 3b-c).

**Fig. 2:**
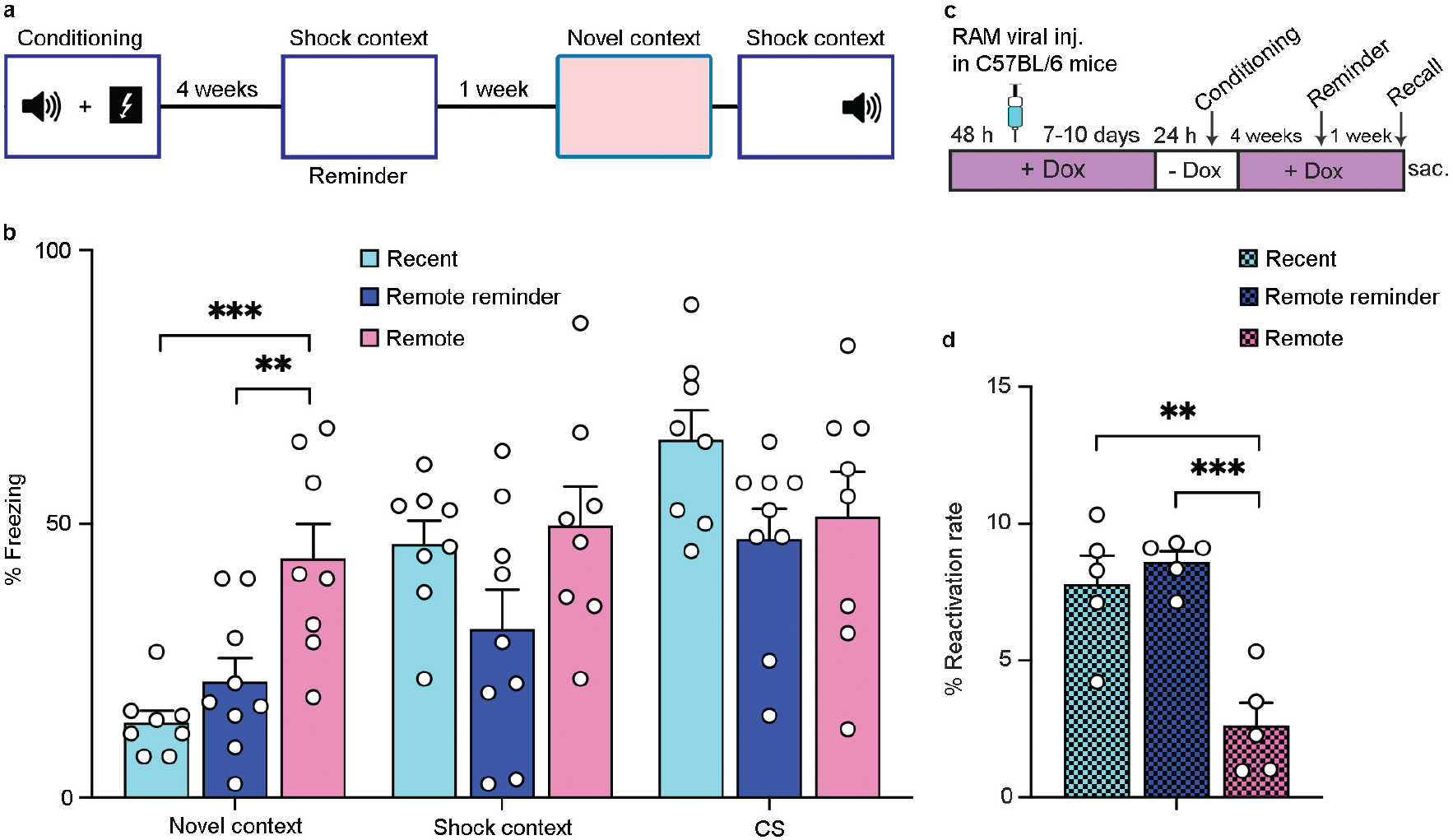
A contextual reminder reinstalls fear engram activation in the dDG and prevents remote memory generalization. **a** Schematic of the remote fear conditioning paradigm with a reminder in the shock-context one week before the recall. **b** Mice with remote reminder (*n* = 9) show significantly decreased freezing behavior during the novel-context exposure than mice without a reminder (*n* = 8). Their freezing behavior in the novel context is comparable to the recent memory group (*n* = 8). Behavior in the shock-context and during conditional tones is not different between groups. **c** Schematic of the RAM system for engram labeling with reminder session. **d** The % reactivation of engram cells in the shock-context after the remote reminder is significantly increased compared to the remote group and reaches levels of the recent group (*n* = 5 each). Data are mean + SEM. Statistics were performed using one-way ANOVA followed by Tukey’s multiple comparison test in **b** and **d**. \*\*\**p*<0.001, \*\**p*<0.005.

To interrogate whether the reduced engram reactivation at a remote time and/or its normalization via a reminder session is potentially due to a dynamic alteration in the circuit excitability of the dDG and its downstream target dCA3, we measured perforant path (PP)-induced field potential signatures in dorsal hippocampal slices obtained at corresponding time-windows. Interestingly, we found no evidence for altered excitability in the dDG (Supplementary Fig. 3d, *F*_2.74_ = 1.117, *p* = 0.333) and mossy fiber (MF)-to-CA3 synapse (Supplementary Fig. 3e, *F*_2.97_ = 0.0435, *p* = 0.957) indicating that the observed engram activation patterns might be rather associated with altered neuromodulation in the dDG-CA3 circuit.

### Remote fear generalization is associated with decreased activity of the NPYergic dDG circuit

Based on our previous observation that fear memory specificity is governed by NPYergic neuromodulation in the dDG during conditioning (27), we next explored the possible role of this circuit in remote memory generalization and reasoned that a change in NPY could be the underlying cause. To test this hypothesis, we first determined the density of NPY immunopositive cells in the dDG in brain sections obtained from the fear-conditioned mice at the recent, remote-reminder, and remote times. Interestingly, we found a significant decrease in the NPY^+^ cell number in the dDG at the remote compared to the recent and remote-reminder group. Of note, NPY^+^ cell numbers were recovered after the reminder session. (Fig. 3a-c, *F*_3.19_ = 7.83, *p* = 0.001; post hoc: recent vs. remote-reminder: *p* = 0.002; recent vs. remote: *p* = 0.002). However, the NPY peptide levels are significantly increased only in the recent group compared to other groups (Fig. 3d, *F*_3.19_ = 7.83, *p* = 0.001; post hoc: recent vs. remote-reminder: *p* = 0.002; recent vs. remote: *p* = 0.002) whereas the NPY plasma levels were not changed between groups at the same times (Supplementary Fig. 4a). These data suggest that the time-dependent decrease in the NPYergic signaling in the dDG circuit entails novel-context generalization in our fear-conditioning paradigm.

**Fig. 3:**
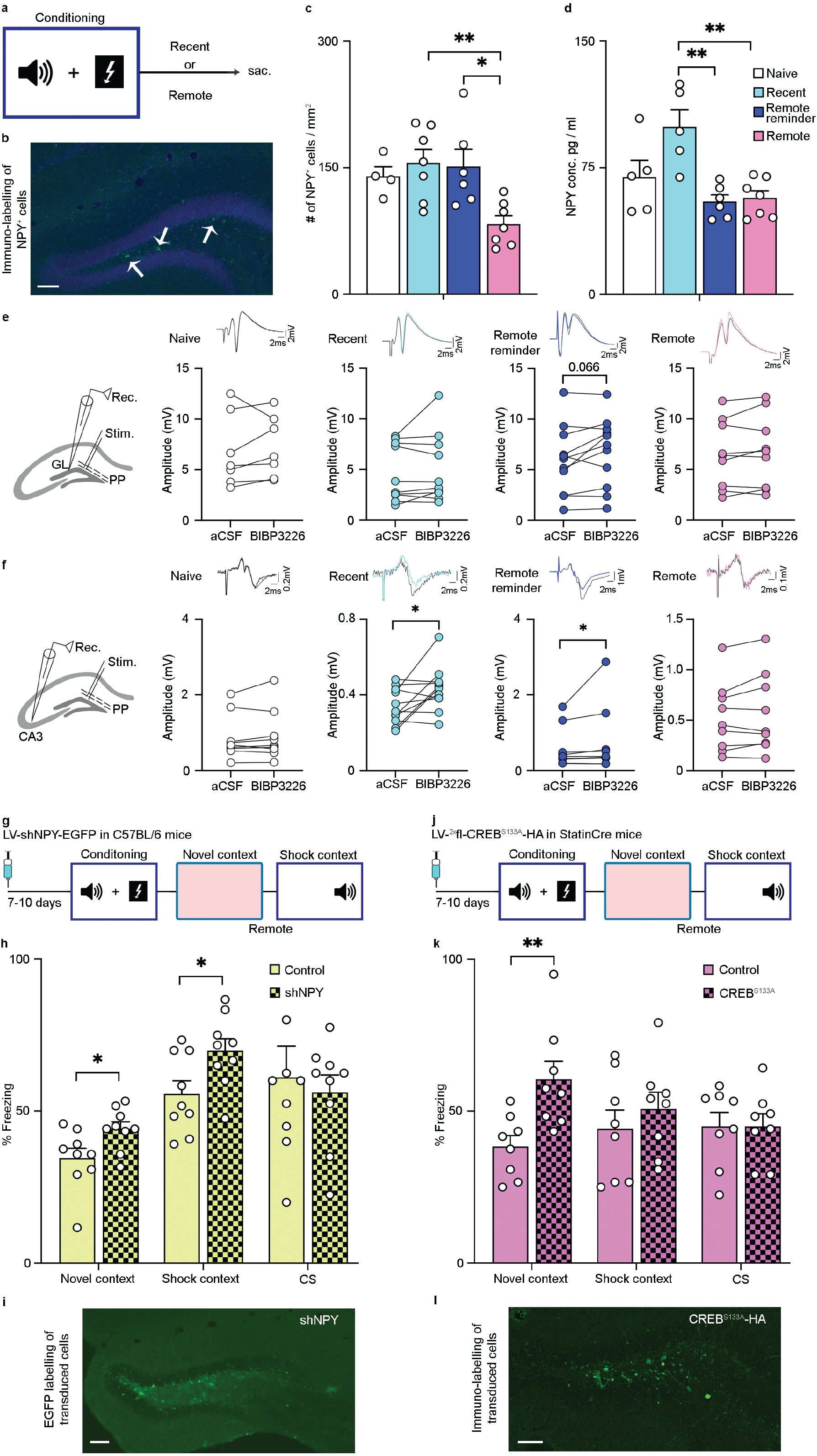
NPY cells in the dDG are involved in remote fear generalization. **a** Schematic overview of conditioning and tissue preparation; mice were sacrificed at time points resembling the recent or remote test sessions but without context exposure. **b** A representative microscopic image shows immune-labeling of NPY^+^ cells in the dDG (arrows). Scale bar, 100 μm. **c** Remote reminder (*n* = 6) increases the number of NPY^+^ cells compared to the remote group (*n* = 7) and reinstates levels comparable to the recent memory (*n* = 7) and naïve groups (*n* = 4). **d** However, the remote reminder does not increase the concentration of NPY peptide in the dDG. **e** Baseline excitability in dorsal dentate gyrus (dDG) granule cell layer (GL) is unaffected by bath-application of NPY-Y1-receptor antagonist BIBP3226 (1μM) in all groups (brain slices-naive: *n* = 7; recent: *n* = 10; remote reminder: *n* = 11; remote: *n* = 9). However, a tendency for increased baseline excitability upon NPY-Y1-receptor antagonism is evident in the remote reminder group. **f** Baseline transmission in the mossy fiber (MF)-dCA3 (perforant-path (PP)- induced disynaptic field responses) is significantly augmented in the recent (*n* = 11 slices) and remote reminder (*n* = 10 slices) group after bath-application of BIBP3226 while no such effect is evident in the naïve (*n* = 9 slices) and the remote (*n* = 9 slices) groups. Representative dDG population spike (PS) responses in **e** and fEPSP responses in **f** before and after BIBP3226 (gray) are shown above the corresponding before-after graphs. The recording schemas for **e** and **f** are illustrated on the left side. **g** Schematic of behavioral experiments with lentiviral NPY knockdown in C57BL/6 mice. **h** shNPY injected animals show a significant increase in the freezing behavior in the novel- and shock-contexts compared to the control injected group. There is no difference between groups upon CS presentation (*n* = 9 each). **i** A representative microscopic image shows the expression of the viral EGFP tag in the hilus. Scale bar, 100 μm. **j** Schematic of behavioral experiments with dominant-negative CREB^S133A^ in StatinCre driver mice. **k** The CREB^S133A^ injected group shows a significant increase in freezing behavior in the novel-context compared to the control injected group. In the shock-context and during CS, freezing behavior remains unchanged between groups (*n* = 8 each). **l** A representative microscopic image of the haemagglutinin (HA) immune-labeling shows viral construct expression in the hilus. Scale bar, 100 μm. Data are mean + SEM. Statistics were performed using one-way ANOVA followed by Tukey’s multiple comparison test in **c** and **d**, and with the Student’s two-tailed paired t-test for normally distributed data or two-tailed Wilcoxon test for non-normal distributed data in **e** and **f**, and with the Student’s two-tailed *t*-test in **h** and **k**. \*\**p*<0.005, \**p*<0.05.

We have previously shown that NPY-Y1 receptor blockade can be used as a sensitive method to interrogate the impact of endogenous NPY on the dDG circuit excitability (27), aligning well with the profound expression of NPY-Y1 receptor in the dDG-CA3 circuit (47). Thus, we measured the modulation of PP-induced DG PS responses and disynaptic MF-mediated fEPSP responses in the dCA3 via NPY-Y1 receptor antagonist BIBP3226 (1 μM). We found no evidence for modulation of dDG excitability (Fig. 3e) via NPY-Y1 receptor antagonism in the slices obtained from the naïve (*t*_6_ = 0.785, *p* = 0.462), recent (*W* = −13.00, *p* = 0.557) or remote group (*t*_8_ = 1.272, *p* = 0.239) but a trend for an increase in the remote-reminder group (*t*_10_ = 2.060, *p* = 0.066). On the other hand, disynaptic MF-dCA3 transmission (Fig. 3f) was significantly increased upon NPY-Y1 receptor antagonism in the slices obtained from recent (*t*_10_ = 2.468, *p* = 0.033) and remote-reminder (*W* = −43.00, *p* = 0.027) groups but not in the slices of naïve (*t*_8_ = 0.927, *p* = 0.381) and remote (*t*_8_ = 1.490, *p* = 0.174) groups. These data indicate that a recent fear conditioning episode leads to a transient augmentation of the NPYergic neuromodulation of the dDG-MF circuit that is reduced at a remote time. Of note, a remote-reminder session seems to reinstate a recent memory-like NPYergic neuromodulation in the dDG-MF circuit.

To further understand the putative role of NPY in controlling remote memory generalization, we knockdown NPY locally in the dDG (33). C57BL/6 mice injected with lentivirus-based shNPY to knockdown NPY or shRandom as control were fear-conditioned and tested for remote memory as described (Fig. 3g). Again, no difference in baseline and post-shock freezing between groups was observed (Supplementary Fig. 4b). However, shNPY significantly increased freezing in the novel-context (*t*_16_ = 2.37, *p* = 0.03) and in the shock-context (*t*_16_ = 2.42, *p* = 0.02) compared to controls without affecting tone memory (Fig. 3g-i). Transcription of the *npy* gene is modulated by transcription factor CREB, and the lentivirus-based conditional expression of dominant-negative CREB (CREB^S133A^) in the HIPP cells results in reduced NPY mRNA expression levels (27). Thus, to examine the chronic contribution of CREB activity in the NPY dependent generalization, we injected CREB^S133A^ into the dDG of StatinCre driver mice (Fig. 3j). As expected, CREB^S133A^ injections significantly increased freezing behavior compared to the controls in the novel-context (*t*_14_ = 3.14, *p* = 0.007), while memory in the shock-context and during tones was not changed (Fig. 3k-l). There was no difference in baseline and post-shock freezing between groups (Supplementary Fig. 4c). This CREB^S133A^ manipulation was effective only when done before conditioning, as we did not observe any difference in mice injected with CREB^S133A^ after fear-training (Supplementary Fig. 4d). However, neither shNPY nor CREB^S133A^ injections changed the number of cFos^+^ cells in the dDG after novel-context exposure (Supplementary Fig. 4e-f). These experiments indicate that chronic reduction of NPY signaling leads to over-generalization of remote fear, but without altering the magnitude of engram reactivation by the novel context.

The reconstitution of basal recent memory-like NPY conditions in the dDG after remote-reminder led us to further test the potentially permissive role of reduced NPY levels for generalization. However, reducing NPY expression by shNPY and CREB^S133A^ manipulation of HIPP cells did not reduce the effectiveness of the reminder session to attenuate the generalization of the novel context or performance in the reminder session itself (Supplementary Fig. 5a-b). Together these data indicate that while NPY levels in dDG are recovered by a fear reminder and associated engram reactivation, NPY itself is not required for the effects of the reminder session.

### Chemogenetic manipulation of the NPYergic circuit in the dDG at the time of fear conditioning determines remote generalization

It has been shown that the acute chemogenetic inhibition of HIPP cells during the acquisition of fear memory is crucial to maintain the specificity of recent memory and respective ensemble activation in the dDG (26–27). To examine acute HIPP cell functioning in remote memory during acquisition, consolidation, and recall, we used designer receptors exclusively activated by designer drug (DREADD) in StatinCre driver mice. We used a Cre-dependent AAV vector expressing hM4Di that inhibits target cell upon clozapine-N-oxide (CNO) drug and vectors expressing only mCherry as control.

Silencing HIPP cells during acquisition resulted in a significant decrease in freezing behavior during remote recall in the novel-context compared to the controls (*t*_18_ = 2.12, *p* = 0.04) with unchanged freezing in the shock-context and during tones (Fig. 4a-c). Importantly, engram labeling at the remote recall under this manipulation was unchanged (Fig. 4d and Supplementary Fig. 6b-c). By contrast, the silencing of HIPP cells immediately and 120 min after conditioning targeting consolidation phase resulted in increased freezing behavior during recall in the novel-context compared to the controls (*t*_13_ = 2.20, *p* = 0.04), with freezing behavior in the shock-context remaining unaltered and freezing behavior during tones significantly decreased (Fig. 4e-f). These data indicate that HIPP cell activity during and shortly after fear memory acquisition determines later generalization. By contrast, neither the silencing of the HIPP cells nor their activation with hM3Dq before remote recall in the novel-context altered freezing behavior in the novel-context, shock-context, or during tones (Fig. 4g-h). The baseline and post-shock freezing behavior in all these groups were unchanged (Supplementary Fig. 6a, d-e). Similar to lentiviral experiments with remote-reminder, neither acute activation nor inhibition of HIPP cells before the reminder session altered freezing behavior during later recall (Supplementary Fig. 6f).

**Fig. 4:**
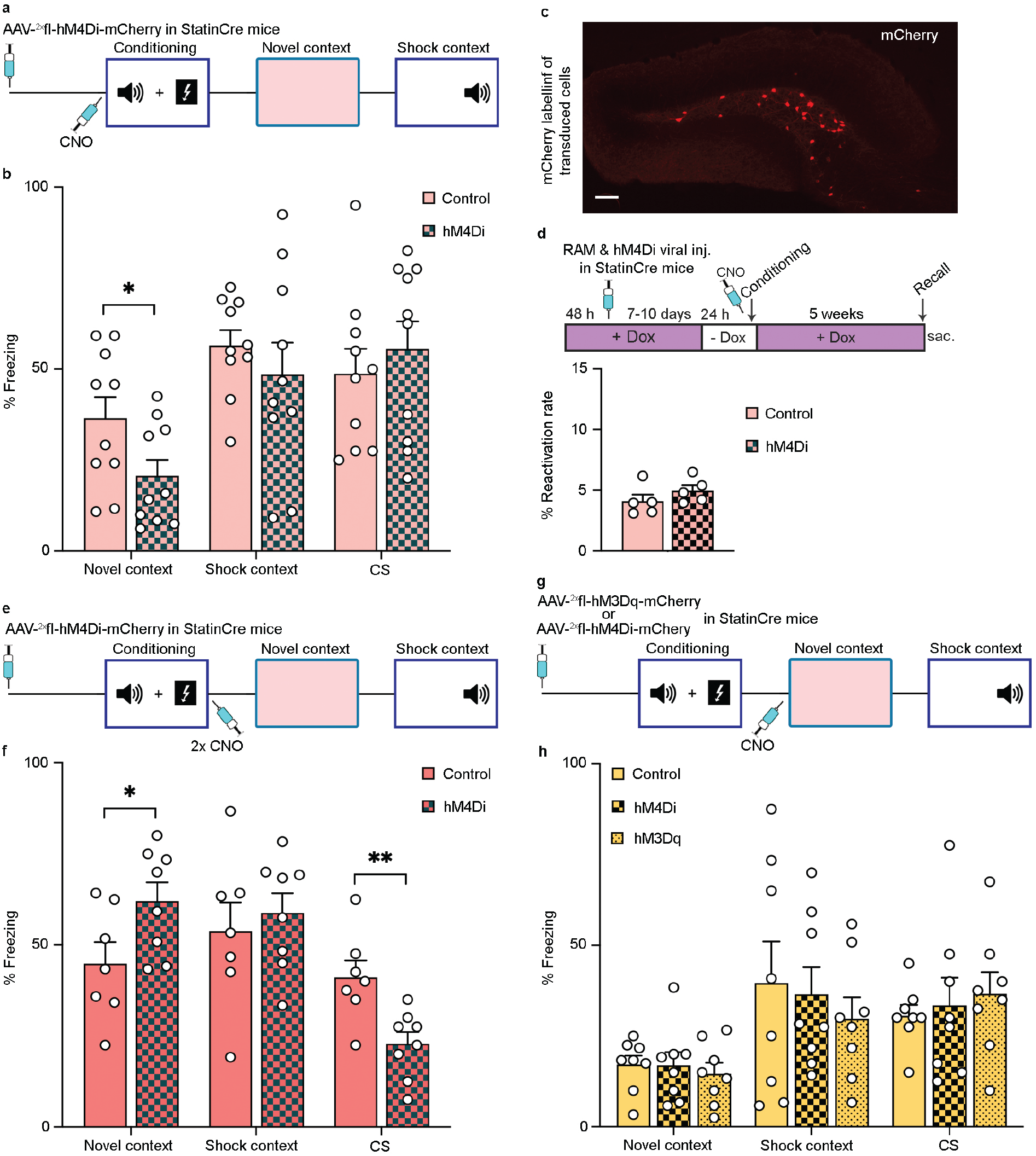
Chemogenetic control of NPYergic HIPP cell activity reveals a role in the acquisition and early consolidation. **a** Schematic of the hM4Di injection paradigm targeting acquisition of fear conditioning. HIPP cells are silenced with i.p. injection of CNO one hour before conditioning. **b** Silencing HIPP cells in hM4Di injected StatinCre mice during acquisition results in a significant decrease in remote memory generalization to the novel context. Freezing behavior is not different between groups in the shock-context or during CS presentation (*n* = 10 each). **c** A representative microscopic image shows a viral expression of the mCherry tag in the dDG. Scale bar, 100 μm. **d** (upper) Schematic of the RAM system for engram labeling and hM4Di injection with i.p CNO before conditioning. (lower) The reactivation of RAM-labelled dDG engram cells during remote generalization is not changed compared to the control group (*n* = 5 each). **e** Schematic of the hM4Di injection paradigm targeting early consolidation of fear memory. HIPP cells are silenced with two CNO injections immediately and 2 h after conditioning. **f** HIPP cell silencing during consolidation significantly increases fear generalization to the novel-context compared to the controls (*n* = 8 each). Freezing behavior is not changed in the shock-context, but significantly decreased during CS presentation compared to the control group. **g** Schematic of hM4Di/hM3Dq injections paradigms to target fear memory recall during the remote generalization session. HIPP cells are silenced/activated with CNO one hour before the novel context exposure. **h** Neither silencing (*n* = 7) nor activation (*n* =8) of HIPP cells before recall changes freezing behavior compared to the control group (*n* = 8) in the novel-context, shock-context, or during CS presentation. Data are mean + SEM. Statistics were performed using Student’s two-tailed *t*-test. \*\**p*<0.005, \**p*<0.05.

## Discussion

This study demonstrates that recall of remote fear memory by reinstating a recent memory-like NPY function and engram activation in the dDG counteracts the spontaneous generalization of remote fear memory. Our data indicate that NPYergic transmission is not only recruited following fear memory recall to modulate DG-CA3 signaling but, critically, already at the time of memory acquisition and early consolidation determines the degree of later generalization. Our findings strongly support the role of NPYergic interneurons of the dentate gyrus in controlling the strength and specificity of fear memories and preventing stress-induced psychopathology.

With time, fear memories tend to become less specific and generalized. While this may help organisms adapt to gradual alterations in an ever-changing environment, an over-generalization of fear is maladaptive and a central symptomatic dimension of post-traumatic stress disorder (PTSD) and anxiety-related disorders (1, 5–7, 48–50). In this study, we employed a Pavlovian fear conditioning protocol that allows assessing memory to an auditory cue and the background context at both recent and remote time points. It thus resembles real-life trauma situations by combining conditioning to an explicit threat with a predictive environmental setting. Previously we and others showed that the salience of the background context information is reflected in the size of the cellular engram activated in the dDG and controlled by NPYergic transmission originating from dDG HIPP cells (26–27). We now could confirm the gradual increase of fear memory generalization to a novel-context over time (Fig. 1) that is accompanied by a loss of engram activation and a reduction of NPYergic function in the dDG at the remote time point. Remote memory generalization was observed regardless of whether mice were single caged or group-housed and whether paired or unpaired auditory conditioning was applied (Supplementary Fig. 1b-c).

Reactivation of memories has been shown to induce a reconsolidation process that renders long-term consolidated memory labile for additional updating and restores memory specificity (51–53). Similarly, reactivation of remote memory has been proposed to update the original hippocampal trace and thereby reduce generalization (39, 54). In fact, we found that presenting a contextual reminder session one week before testing remote memory test not only prevents generalization but also leads to enhanced recruitment of fear memory engram cells in the dDG during exposure to a novel context (Fig. 2). Granule cells are incorporated into fear memory engrams (18), and stimulating these pre-formed engrams in a safe environment results in fear memory retrieval (19). These engrams also recruit local interneurons in the dDG, including NPY^+^ cells, to inhibit non-engram cells from supporting pattern separation and memory specificity (26–28). During recent memory, mice show increased fear in the shock-context compared to the novel-context indicating intact memory specificity, which corresponds to a higher number of activated engram cells in our RAM labeling experiment. On the other hand, during remote memory, the number of engram cells activation is decreased, as observed by others (55), without a change in the overall activity of granule cells. This suggests that potentially during systems consolidation of memory, the accessibility of dDG engram cells is reduced, and the fear expression is carried out by the cortical regions, especially the anterior cingulate cortex (7, 38). Presumably, this activation of dDG engram cells reflects a recapitulation of previously acquired memory, which according to the dentate gyrus’ function as a pattern separator, can be utilized to discriminate the novel-context and thus prevent generalization. The remote fear memory after a reminder thus appears to be “refreshed”, resembling the situation during recent fear memory recall.

In line with this interpretation, the reminder session induced a recovery of recent memory-like physiological and neurochemical parameters in the dDG, including the number of NPY-positive interneurons and NPYergic control of excitability in the dDG-CA3 pathway. It is plausible that a decrease in the NPY^+^ cell number and NPY peptide in the dDG at the remote time point (Fig. 3c-d) may result in reduced inhibition of dDG granule cells or mossy fibers and thus permit generalization. We observed no effect of NPY-Y1-receptor blockade on PP-induced excitability in the dDG hippocampal slices, in line with previous observations (27, 56). By contrast, we found that NPY-Y1 receptor blockade augmented disynaptic MF-mediated fEPSP in recent and remote-reminder groups. This finding aligns well with a previous study assessing PP-induced responses in the dCA3 *in vivo* (57). Thus, the NPYergic system in the dDG-MF circuit appears to be transiently recruited following fear-conditioning. Reinstating this function by the reminder session and the observed engram activation may help restore pattern separation functions of the dDG and thereby improve context discrimination.

Accordingly, knockdown of NPY in dDG and suppression of activity-dependent NPY expression in dDG HIPP cells with dominant-negative CREB^S133A^ led to an over-generalization of fear. However, the CREB^S133A^ manipulation was not effective when carried out after memory consolidation (Supplementary Fig. 4d). This prompted us to test further the time-dependent role of HIPP cells with chemogenetic silencing, which suggested that activation of these interneurons during memory acquisition contributes to remote memory generalization. This effect is not simply explained by the previously identified role in encoding background context salience (27). In fact, no alteration of context memory strength could be observed in the remote time point. Moreover, HIPP cells activation and silencing had no effect during generalization testing, in agreement with our previous findings on memory retrieval. However, HIPP cell silencing during fear memory consolidation enhanced memory generalization, similar to the NPY knockdown and CREB^S133A^ experiment. This suggests that post-training activity in the dDG is essential to limit the remote generalization of conditioned fear, and deficits in the consolidation-related function of these cells may support pathological generalization.

NPY in the brain has anxiolytic properties and attenuates fear-conditioned memories as a control mechanism to cope with inappropriate or exaggerated fear after stressful and traumatic events (30–31). Indeed, several lines of evidence suggest that NPY may be a resilience factor in PTSD, as, e.g., low NPY levels are associated with the presence of intrusion symptoms (58) and high NPY is predictive of PTSD symptom improvement and coping following a traumatic event (29). Treatment strategies against the pathophysiology of PTSD include fear extinction and fear reminder mediated fear inhibition, which are widely used that possibly recruit hippocampal processes (5, 59–60). Our data identify a specific mechanism of fear memory generalization under NPYergic control that may be targeted to relieve memory generalization in these conditions.

## Acknowledgements

We are grateful to F. Webers and S. Stork for expert technical assistance and A. Bohnsted and D. Al-Chakmachi for excellent animal care. We are thankful to F. A. Manjili for helping with immunostainings. This work is supported by the European Regional Development Fund (ERDF), Project Center for Behavioral Brain Sciences (CBBS), FKZ: ZS/2016/04/78113 to S.A.R and the German Research Foundation (362321501/RTG 2413 SynAGE TP10 to O.S., CRC1436 project Z01 to O.S. and M.R.K.).

## Author contributions

S.A.R and O.S conceptualized the research; S.A.R carried out experiments and analyzed data. K.K carried out field recording experiments with inputs from G.Ç; M.D.-Á carried out ELISA experiments; Y.E.D developed lentiviral constructs. M.R.K developed RAM vectors. S.A.R wrote the original draft of the manuscript; S.A.R and O.S reviewed and edited the manuscript with input from all authors.

