## Supplementary information for "Fear engrams and NPYergic circuit in the dorsal dentate gyrus determine remote fear memory generalization"

Running title: NPY in the dDG determines remote generalization

Syed Ahsan Raza[1, 2*], Katharina Klinger[1ε], Miguel del Ángel[1ε], Yunus Emre Demiray[1], Gürsel Çalışkan[1, 2], Michael R. Kreutz[2, 3, 4] & Oliver Stork[1, 2*]

**Affiliations:**

[1]Department of Genetics and Molecular Neurobiology, Institute of Biology, Otto-von-Guericke University, 39120 Magdeburg, Germany.

[2]Center for Behavioral Brain Sciences, Magdeburg, Germany.

[3]RG Neuroplasticity, Leibniz Institute for Neurobiology, 39118 Magdeburg, Germany

[4]Leibniz Group 'Dendritic Organelles and Synaptic Function,' University Medical Center Hamburg-Eppendorf, Center for Molecular Neurobiology, ZMNH, 20251 Hamburg, Germany

[ε]These authors contributed equally to this work.

[*]Corresponding authors

Address: Department of Genetics and Molecular Neurobiology, Institute of Biology, Otto-von-Guericke University Magdeburg, Leipziger Str. 44, Haus 91, 39120 Magdeburg, Germany.

Supplementary Figures, Raza et al

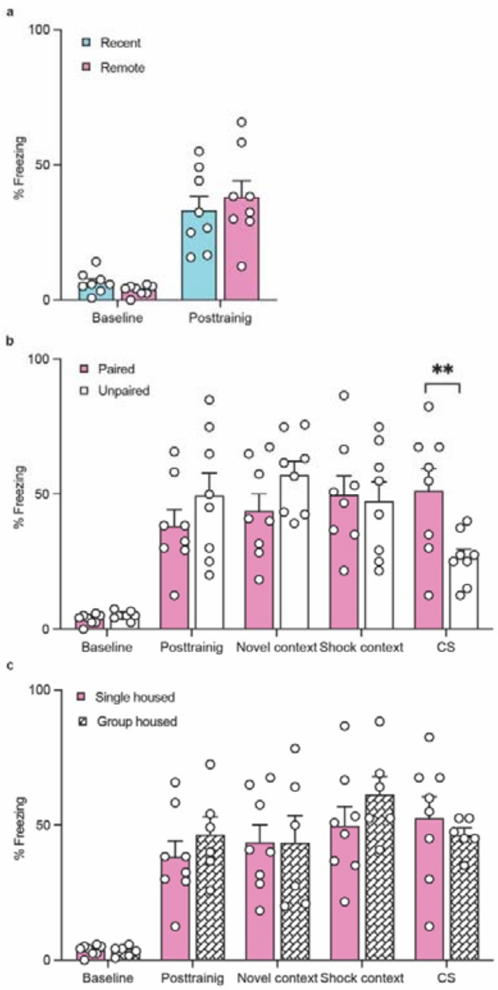


**Supplementary Figure 1: Additional data related to Fig. 1. a** Recent and remote memory groups show similar freezing behavior during baseline and the immediate post-shock interval ($n$ = 8 each). **b** The freezing behavior after paired (CS-US) and unpaired conditioning are similar in the novel-context indicating equal remote memory generalization. As expected, the freezing behavior during CS presentation is significantly higher in paired conditioning than unpaired conditioning, suggesting the salience value is loaded more strongly onto the conditional stimulus in paired conditioning ($n$ = 8 each; $t_{14}$ = 2.80, $p$ = 0.014). **c** Freezing behavior is not different between animals that were group-housed ($n$ = 6) and animals that were single housed ($n$ = 8) after fear conditioning until recall. Data are mean + SEM. Statistics were performed using Student´s two-tailed *t*-test. *$p$<0.05

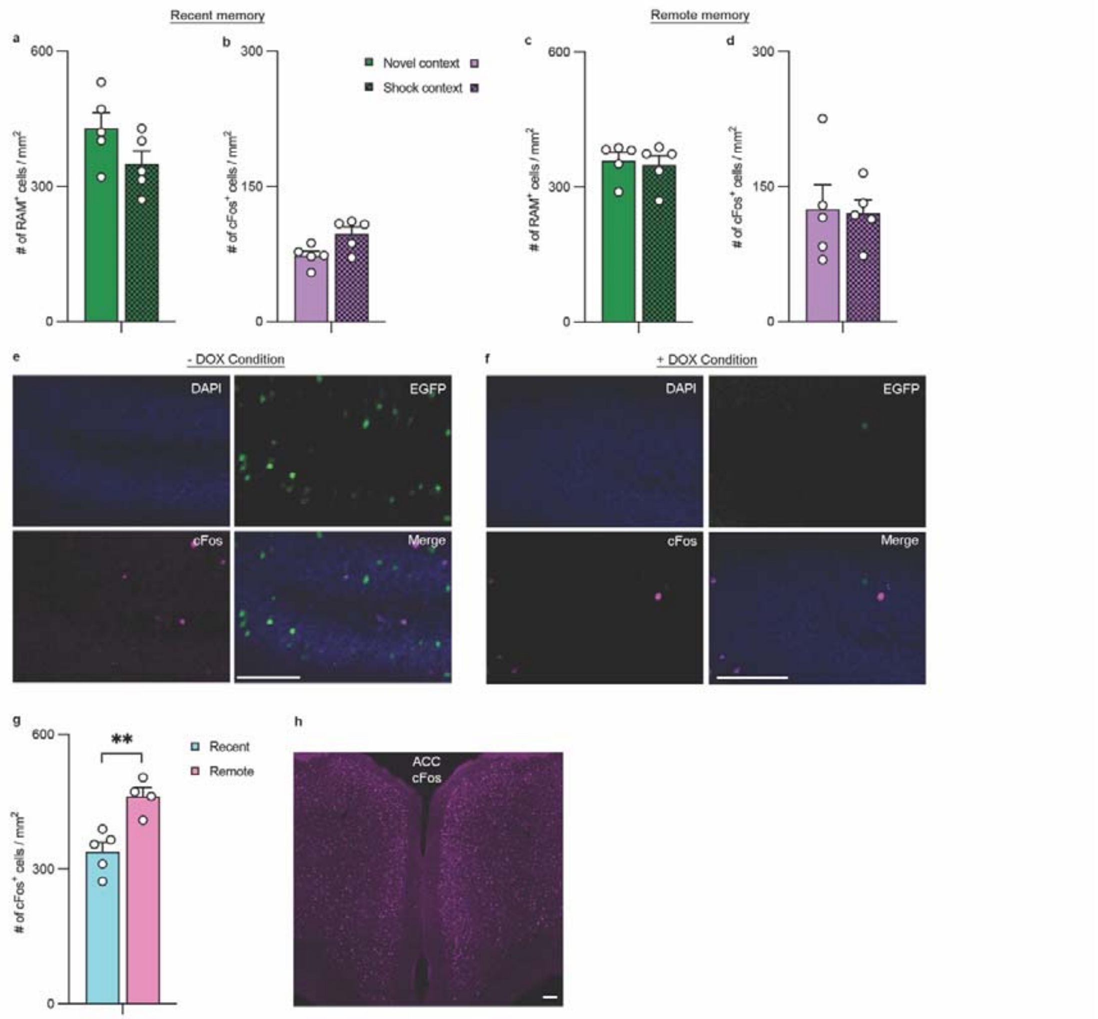


**Supplementary Figure 2: Additional data related to Fig. 1. a** The total number of RAM$^+$ cells labeled with the AAV-RAM-EGFP viral vector during conditioning is not changed after novel vs. shock-context exposure in recent memory recall (*n* = 5 each). **b** The total number of cFos$^+$ cells in dDG is also not different between these groups. Similarly, **c** RAM$^+$ cell number and **d** cFos$^+$ cell number is not different between novel vs. shock-context exposure in remote memory (*n* = 5 each). **e** Representative microscopic images showing expression of EGFP from the RAM construct after conditioning under -Dox chow, whereas in **f**, conditioning under +Dox chow fails to induce the RAM$^+$ marker. DAPI (blue), RAM-EGFP$^+$ (green), cFos$^+$ (magenta) and merge. Scale bar, 100 µm. **g** Comparing cFos$^+$ cell number in the anterior cingulate cortex (ACC) between recent (*n* = 5) and remote (*n* = 4) groups shows higher cFos$^+$ cell number after novel-context exposure in the remote group, which is in line with previous reports (37-38). **h** A microscopic image shows cFos expression in the ACC. Scale bar, 100 µm. Data are mean + SEM. Statistics were performed using Student´s two-tailed *t*-test. $^{**}p<0.005$

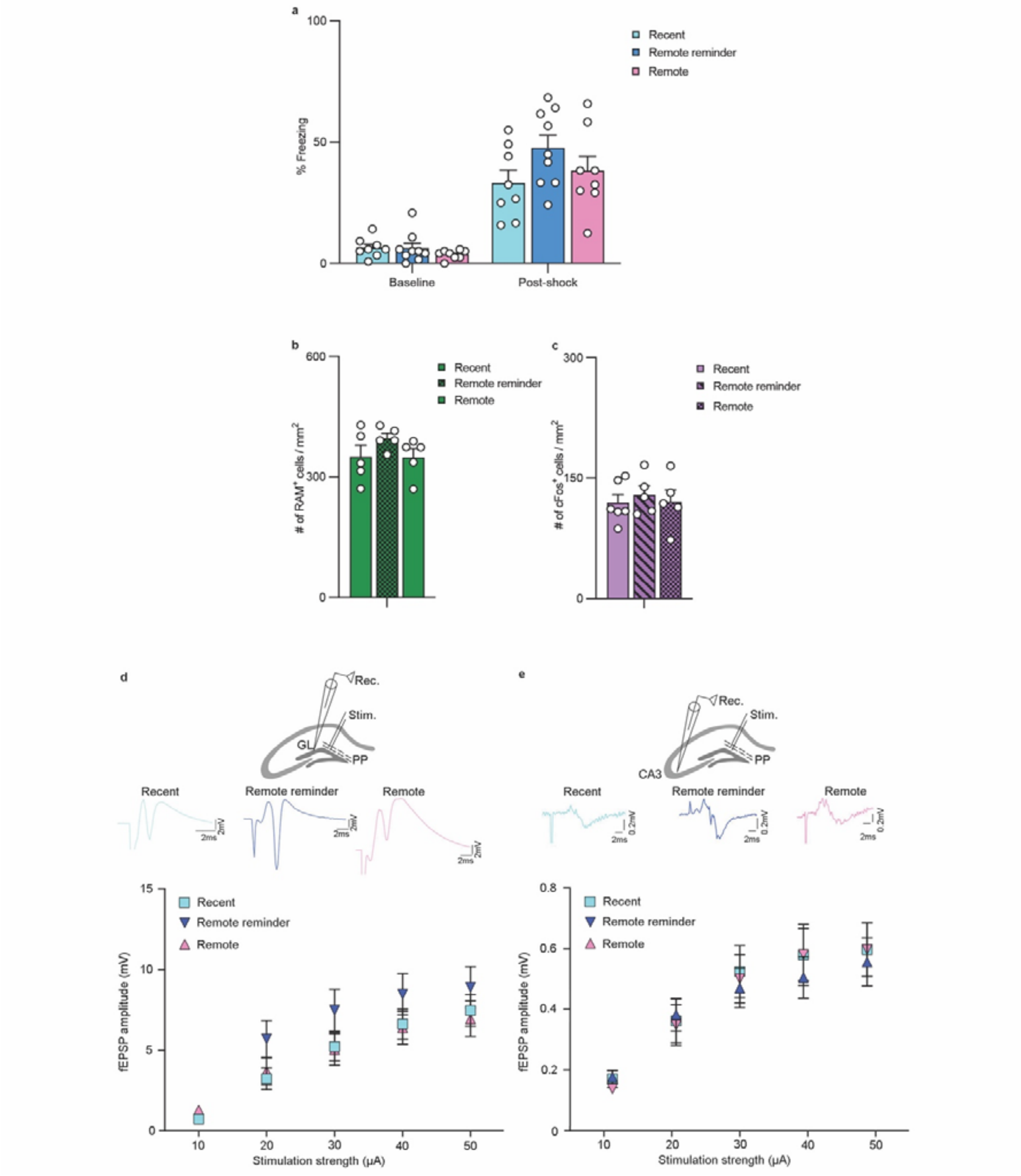


**Supplementary Figure 3: Additional data related to Fig. 2. a** Freezing behavior during baseline and the immediate post-shock phase is not different between recent (*n* = 8), remote reminder (*n* = 9), and remote groups (*n* = 8). **b** The total number of $RAM^+$ cells and **c** $cFos^+$ cells is also not different between the recent, remote reminder, and remote groups (*n* = 5 each). **d** Input / output (I/O) curves of perforant path (PP)-induced population spike (PS) responses in the dorsal dentate gyrus (dDG) granule cell layer (GL) (brain slices-recent: *n* = 24; remote reminder: *n* = 23; remote *n* = 30) and **e** disynaptic mossy fiber (MF)-mediated field excitatory postsynaptic potential (fEPSP) responses in the dCA3 (brain slices-recent: *n* = 36; remote reminder: *n* = 28; remote *n* = 36) remain similar between groups indicating an unaltered baseline excitability in the dDG-CA3. Representative PP-dDG PS and MF-dCA3 fEPSP traces (at 30 μA) for each group and a recording schema, are plotted above the corresponding (I/O) curve. Data are mean ± SEM.

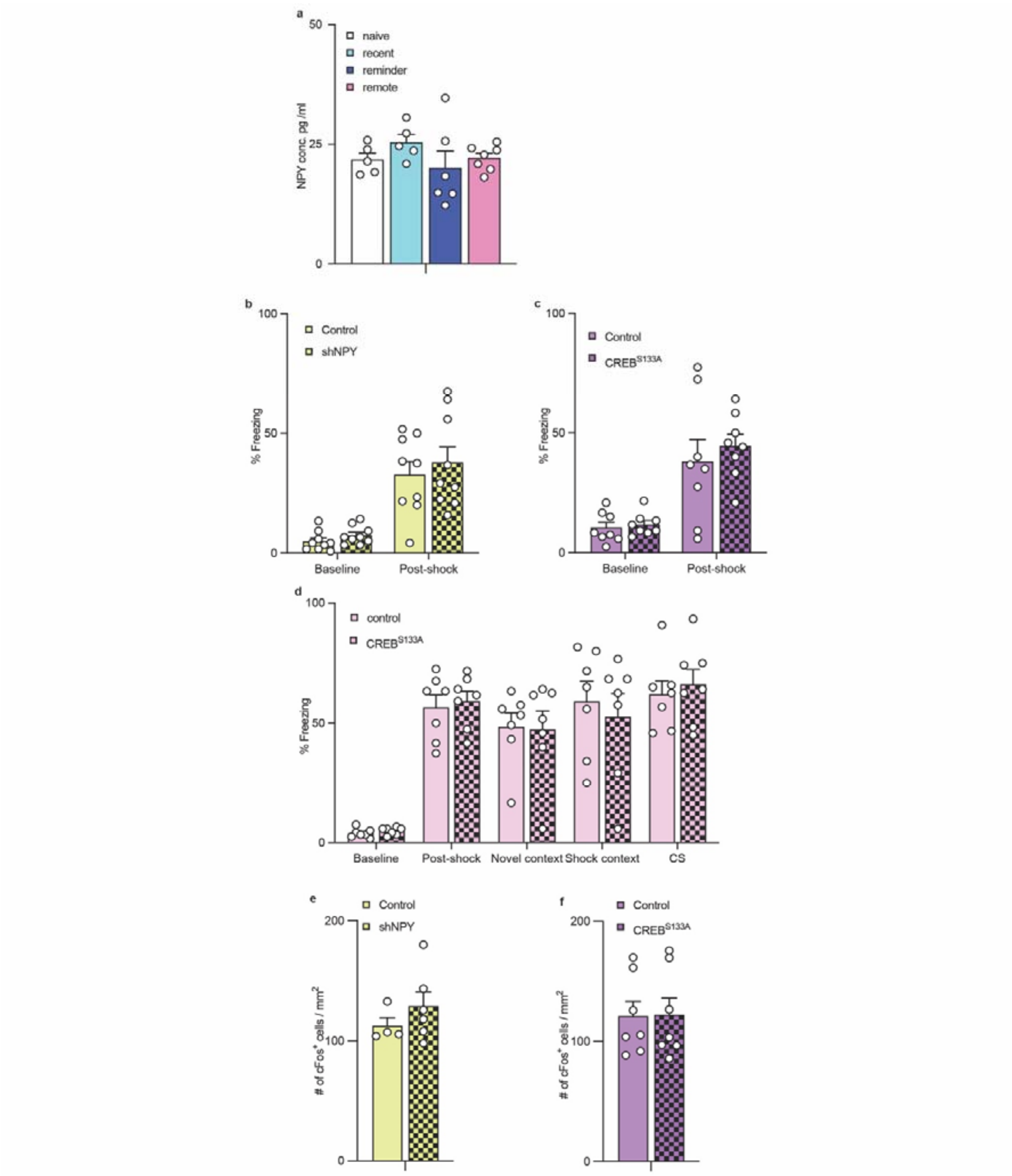


**Supplementary Figure 4: Additional data related to Fig. 3. a** The concentration of NPY peptide in blood plasma is not different between naïve (*n* = 4), recent (*n* = 5), remote reminder (*n* = 6), and remote groups (*n* = 5). **b** Compared to virus injected controls, the baseline freezing behavior and immediate post-shock response are not altered in the shNPY group (*n* = 9 each). **c** Similarly, freezing behavior at these time points is not changed after $CREB^{S133A}$ injection compared to controls (*n* = 8 each). **d** Furthermore, the injection of $CREB^{S133A}$ one week after fear memory training does not change freezing behavior in any phase compared to the group with control virus injection (*n* = 7 each). **e** With another cohort of mice, the number of $cFos^{+}$ cells in brain sections obtained 90 min after novel-context exposure from the shNPY injected group (*n* = 6) is not changed compared to their control group (*n* = 4). **f** The number of $cFos^+$ cells in brain sections obtained from the $CREBS^{S133A}$ injected and control group (*n* = 7) was also not changed. Data are mean + SEM.

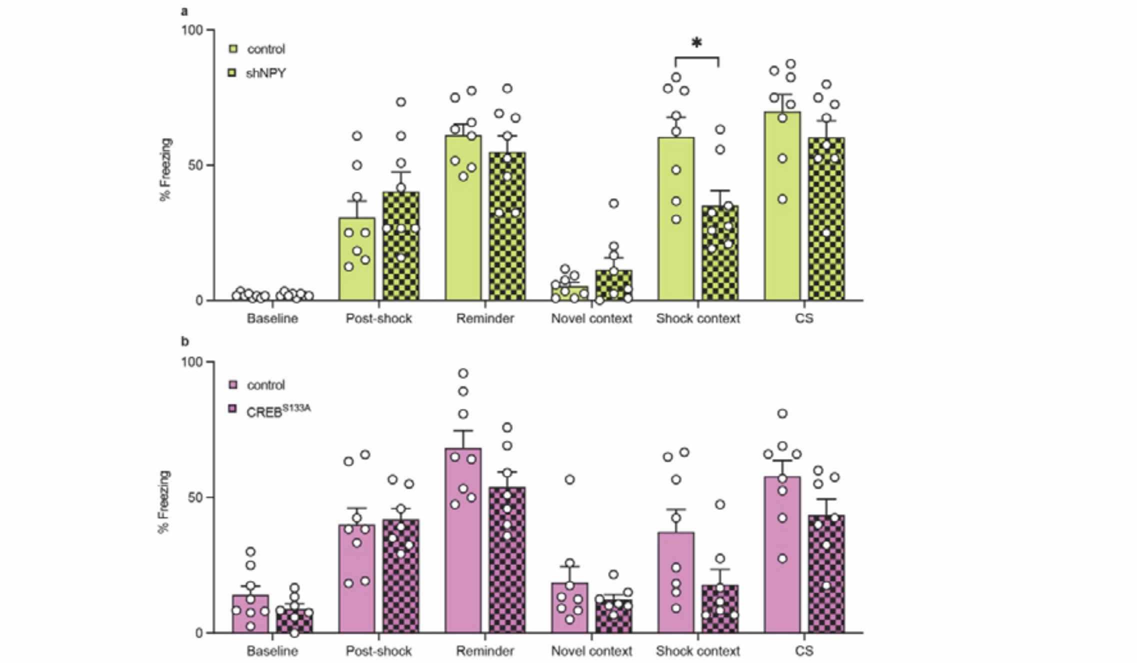


**Supplementary Figure 5: Additional data related to Fig. 3. a** Knockdown of NPY with shNPY expressing viral vectors does not affect baseline freezing behavior or freezing during the immediate post-shock period or the reminder session. However, it significantly decreases the freezing response in the shock-context compared to the control group ($t_{14}$ = 2.80, $p$ = 0.01). The freezing behavior during the novel-context and during the CS presentation is unaltered ($n$ = 8 each). **b** The CREB$^{S133A}$ viral injections ($n$ = 8) do not change freezing behavior in any of the phases of conditioning and testing compared to the controls ($n$ = 7), but a trend is evident towards reduced freezing in the shock-context, similar to that observed upon shNPY mediated knockdown. Data are mean + SEM. Statistics were performed using Student´s two-tailed $t$-test. *$p<0.05$

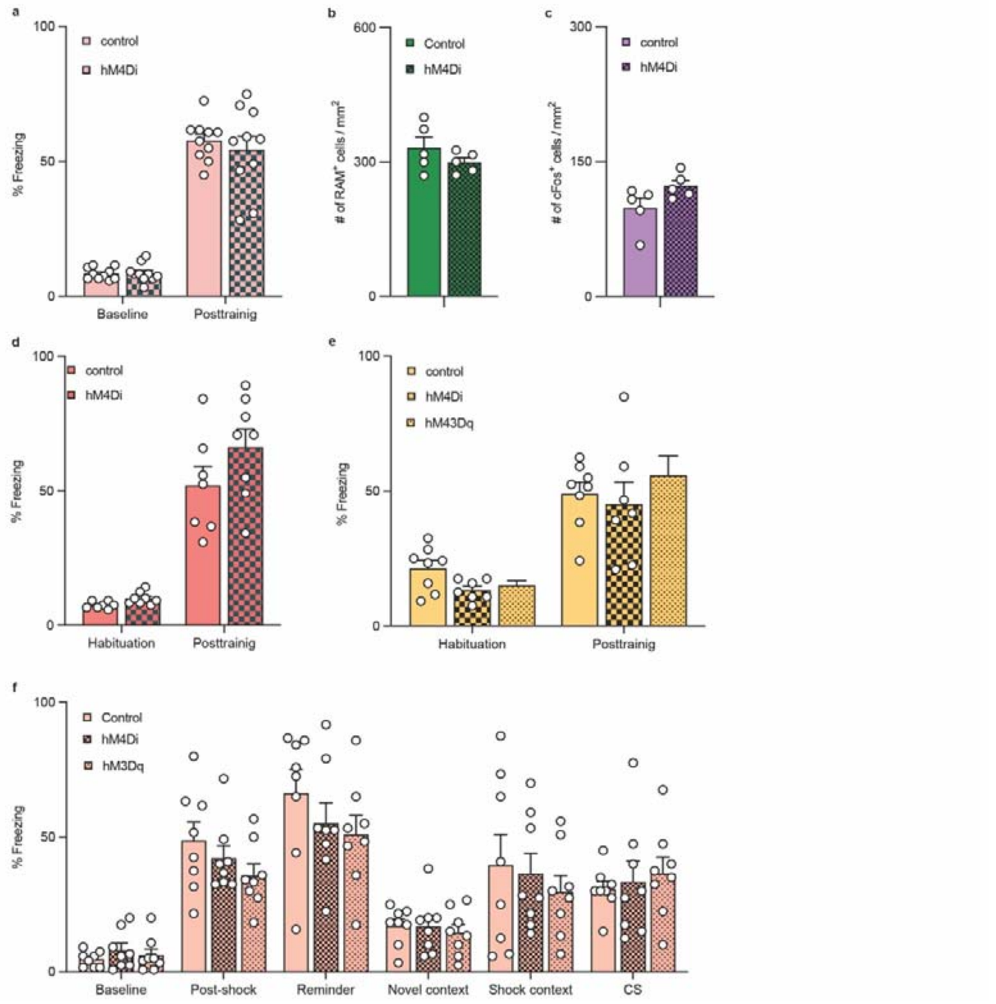


**Supplementary Figure 6: Additional data related to Fig. 4. a** Baseline and immediate post-shock freezing behavior are unaltered in hM4Di expressing animals compared to controls, with CNO injected before fear training (*n* = 10 each). **b** The total number of RAM$^+$ cells or **c** cFos$^+$ cells is not different between hM4Di vs. control groups in the engram labeling experiment (*n* = 5 each). **d** Baseline and immediate post-shock freezing behavior are unaltered in hM4Di expressing mice compared to the control group, which receive CNO after fear training (*n* = 8). **e** Baseline and immediate post-shock freezing behavior are unaltered in hM4Di (*n* = 7) and hM3Dq (*n* = 8) expressing animals compared to controls (*n* = 8), which receive CNO before recall in the remote novel-context. **f** The silencing with hM4Di and activation with hM3Dq of HIPP cells before contextual reminder do not change freezing behavior in any phase of the test protocol (*n* = 8, each). Data are mean + SEM.
